## Supplementary figures and images for "Impact of liver-specific survival motor neuron (SMN) depletion on central nervous system and peripheral tissue pathology"

### Supplementary Figure 1

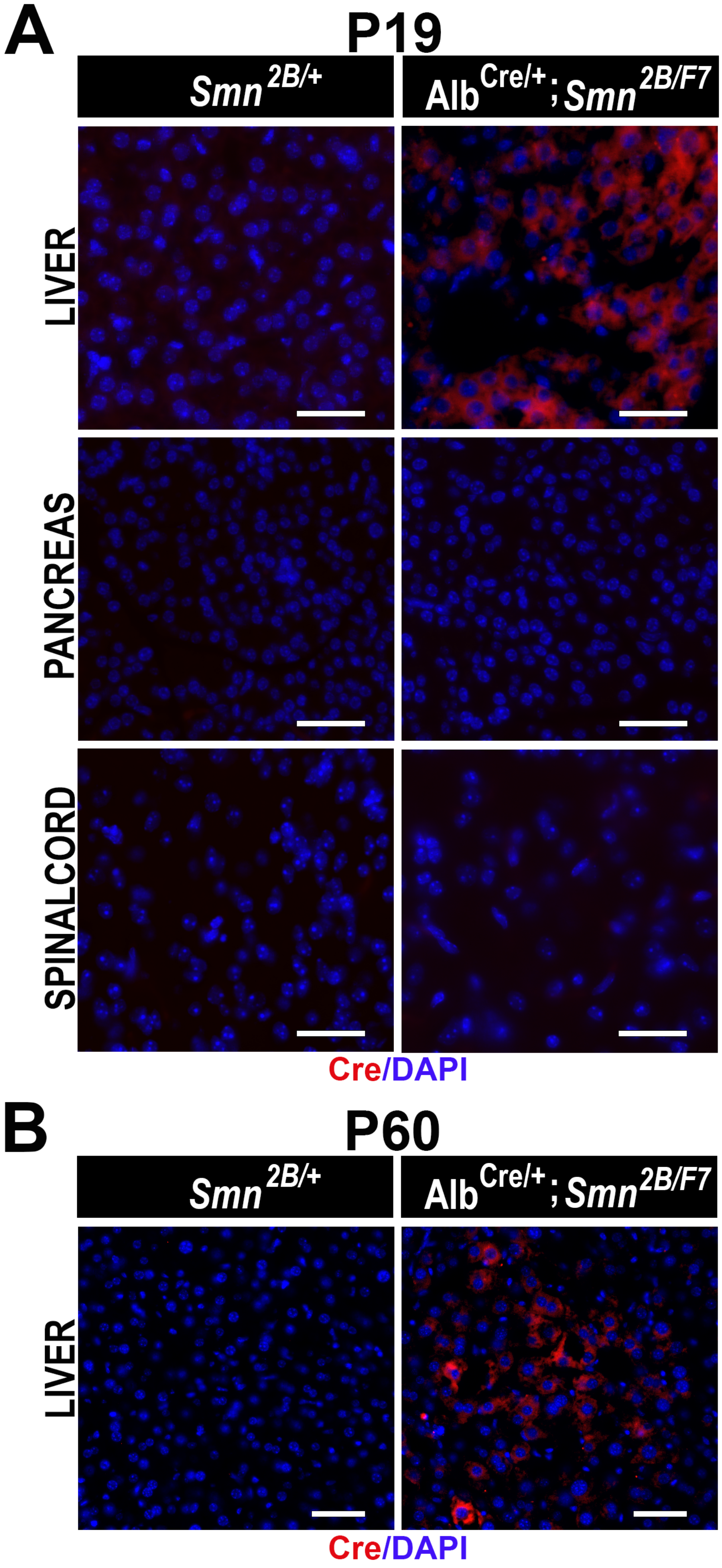
